# Pre-stimulus alpha oscillations encode stimulus-specific visual predictions

**DOI:** 10.1101/2024.03.13.584593

**Authors:** Dorottya Hetenyi, Joost Haarsma, Peter Kok

## Abstract

Predictions of future events have a major impact on how we process sensory signals. However, it remains unclear how the brain keeps predictions online in anticipation of future inputs. Here, we combined magnetoencephalography (MEG) and multivariate decoding techniques to investigate the content of perceptual predictions and their frequency characteristics. Participants were engaged in a shape discrimination task, while auditory cues predicted which specific shape would likely appear. Frequency analysis revealed significant oscillatory fluctuations of predicted shape representations in the pre-stimulus window in the alpha band (10 – 11Hz). Furthermore, we found that this stimulus-specific alpha power was linked to expectation effects on shape discrimination. Our findings demonstrate that sensory predictions are embedded in pre-stimulus alpha oscillations and modulate subsequent perceptual performance, providing a neural mechanism through which the brain deploys perceptual predictions.

## Introduction

Predictions about how the world is structured play an integral role in perception^1–4^. Our prior knowledge forms the basis for predicting future sensory events, which are subsequently integrated with sensory input to form a perceptual experience. While there is a wealth of evidence supporting the idea that the brain deploys predictions to guide perception, the mechanisms through which the brain keeps these predictions online remain largely unclear. One likely candidate for conveying perceptual predictions are neural oscillations^5–8^.

Alpha rhythms (8 – 12Hz) are the predominant oscillations in the awake human brain^9^, yet their functional role is controversial^10,11^. The amplitude and phase of these ongoing oscillations is known to influence performance in visual tasks^12–19^, and have been found to vary with experimental manipulations that target stimulus predictability. Specifically, pre-stimulus alpha oscillations have a similar topography to post-stimulus responses, implying a shared neural substrate in the processing of pre-existing information and external stimuli^20^, and have been shown to predictively encode the position of a moving stimulus^21^. However, whether these oscillations actually convey the contents of perceptual predictions remains unknown.

To test this hypothesis, we employed magnetoencephalography (MEG) combined with multivariate decoding to resolve visual representations with millisecond resolution, and characterise the temporal and frequency characteristics^22,23^ of sensory predictions. Participants were engaged in a shape discrimination task where auditory cues predicted the identity of upcoming abstract shapes. We identified the neural representations of cued sensory predictions prior to stimulus onset and tested whether these sensory predictions had an oscillatory nature, as well as whether the power of such predictive oscillations modulated perceptual performance.

## Results

### Prediction templates oscillate at alpha frequencies

To test whether perceptual predictions are conveyed by oscillations, thirty-two participants performed a challenging visual shape discrimination task (Fig. 1A) while auditory cues predicted the most likely upcoming shape (shape A or D) on 75% of the trials (Fig. 1B). The shape discrimination task was orthogonal to the prediction manipulation (i.e., the cue did not convey any information about whether the two shapes would be identical or different).

**Fig. 1:**
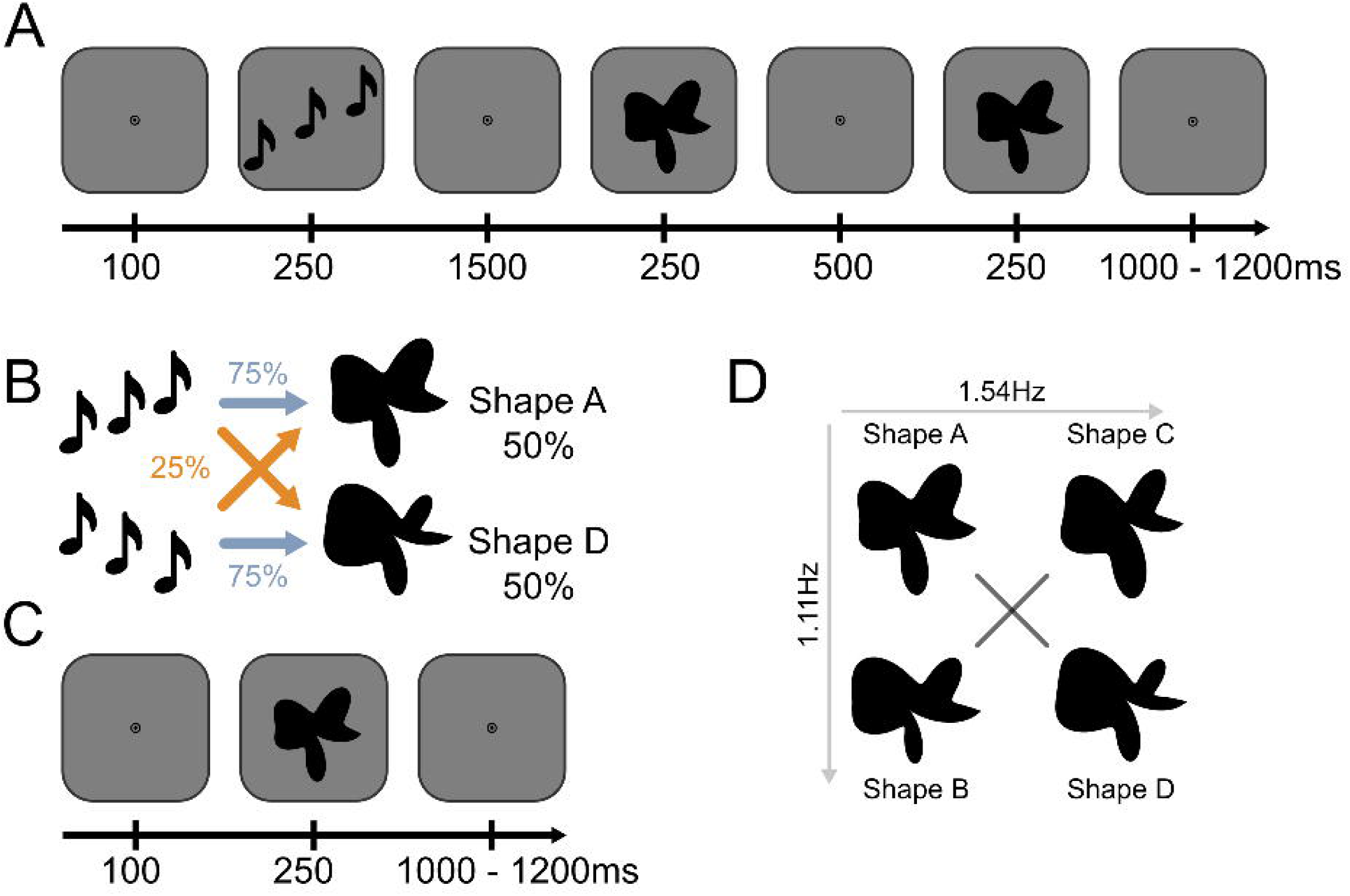
Experimental paradigm. **A:** During prediction runs, an auditory cue preceded the presentation of two consecutive shape stimuli. On each trial, the second shape was either identical to the first or slightly warped with respect to the first along an orthogonal dimension, and participants’ task was to report whether the two shapes were the same or different. **B:** The auditory cue (rising vs. falling tones) predicted whether the first shape on that trial would be shape A or shape D. The cue was valid on 75% of trials, whereas in the other 25% of (invalid) trials the unpredicted shape was presented. **C:** During shape localiser runs no predictive auditory cues were presented and participants performed a fixation diming task. **D:** Four different shapes were presented in the localiser runs, appearing with equal (25%) likelihood. Only shape A and D were presented in the prediction runs. The amplitudes of two RFCs (1.11, and 1.54Hz components) were varied in order to create a two-dimensional shape space, such that shape A vs. D discrimination was orthogonal to shape B vs. C discrimination.

First, we identified shape-specific neural signals using a linear discriminant analysis^24^ (LDA) during separate shape localiser runs (Fig. 1C). Localiser runs consisted of the presentation of four abstract. shapes (Fig. 1D), which were designed to lie on two orthogonal axes of perceptual and neural discriminability (shape A vs. D and shape B vs. C, respectively; see Methods). To test whether the LDA was able to uncover neural representations of the presented shapes, we trained and tested a shape A vs. D decoder within the localiser runs in a cross-validated manner (−100 to 600ms, relative to stimulus onset). We found that the decoder was highly accurate at discriminating the shapes based on the MEG signal. The presented shapes were successfully decoded from 65ms to 450ms (p < 0.001), 465 to 485ms (p = 0.005) and 505ms to 550ms (p = 0.001), peaking at 105ms (Fig. 2A-B). Thus, we could decode abstract shapes during the localiser runs. For all subsequent analyses, decoding traces were averaged over a training window of 70 to 200ms, during which shape decoding peaked (Fig. 2A).

**Fig. 2:**
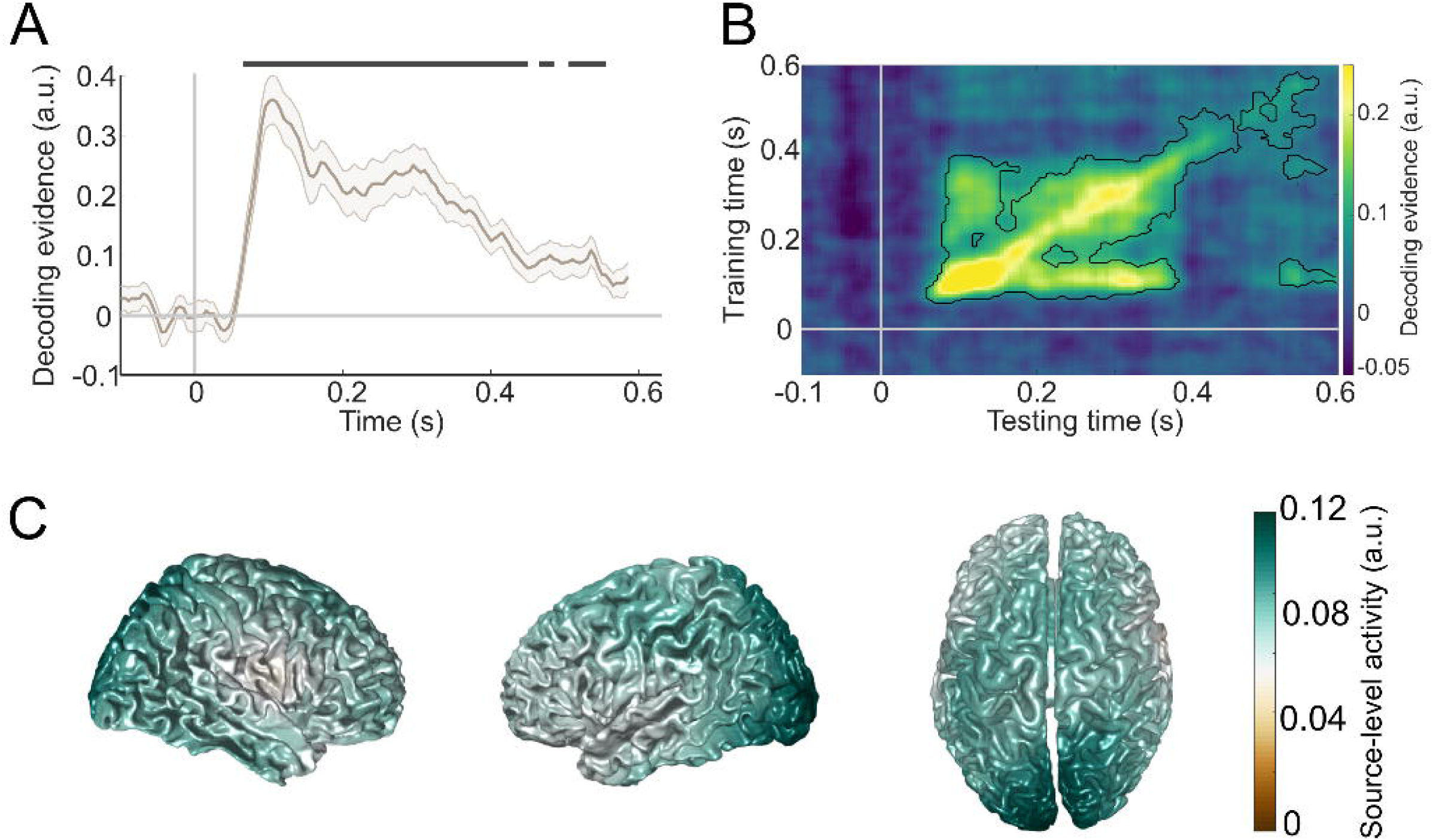
Localiser shape decoding results. **A:** Time course of shape A vs. D decoding. Shapes were successfully decoded from 65ms to 450ms (p < 0.001), 465 to 485ms (p = 0.005) and 505ms to 550ms (p = 0.001), peak at 105ms. Shaded regions indicate SEM. **B:** Temporal generalisation matrix of shape A vs. D decoding, obtained by training decoders on each time point and testing all decoders on all time points. Solid black lines indicate significant clusters (p < 0.05); solid grey lines indicate stimulus onset (t = 0s). **C:** Source localisation of shape A vs. D discrimination during the localiser, training time window of 70 to 200ms post-stimulus, indicating strong occipital activity.

To reveal the cortical sources contributing to these shape representations, we performed source localisation analyses using an linearly constrained minimum variance (LCMV) beamformer^25^ where calculating the difference of the source-localised ERF between shape A and D reflects the contributing brain areas^26^. This revealed strong signals in the occipital lobe, predominantly over the visual cortices (Fig. 2C), in line with the hypothesised sensory nature of these representations.

Next, we used this decoder trained on the localiser to test whether the predictive auditory cues induced oscillatory representations of the predicted abstract shapes (Fig. 3A). To establish the specificity of the neural signals induced by predictions, we created two separate baseline measurements. First, we shuffled the shape labels before training the decoder (N=25 permutations per participant) in order to create a bootstrapped baseline (Baseline 1, Fig. 3B - top). Second, we trained a decoder to distinguish two shapes which were presented in the localiser runs, but not in the main experiment runs (shapes B and C; Fig. 1D). This discrimination was orthogonal to shape A vs. D discrimination, which was confirmed via an absence of generalisation between the two decoders (Fig. S1). The shape B vs. C decoder thus provides a highly specific baseline (Baseline 2, Fig. 3B - bottom), since it was trained to pick up neural representations of highly similar but orthogonal shapes to those that were predicted by the auditory cues.

**Fig. 3:**
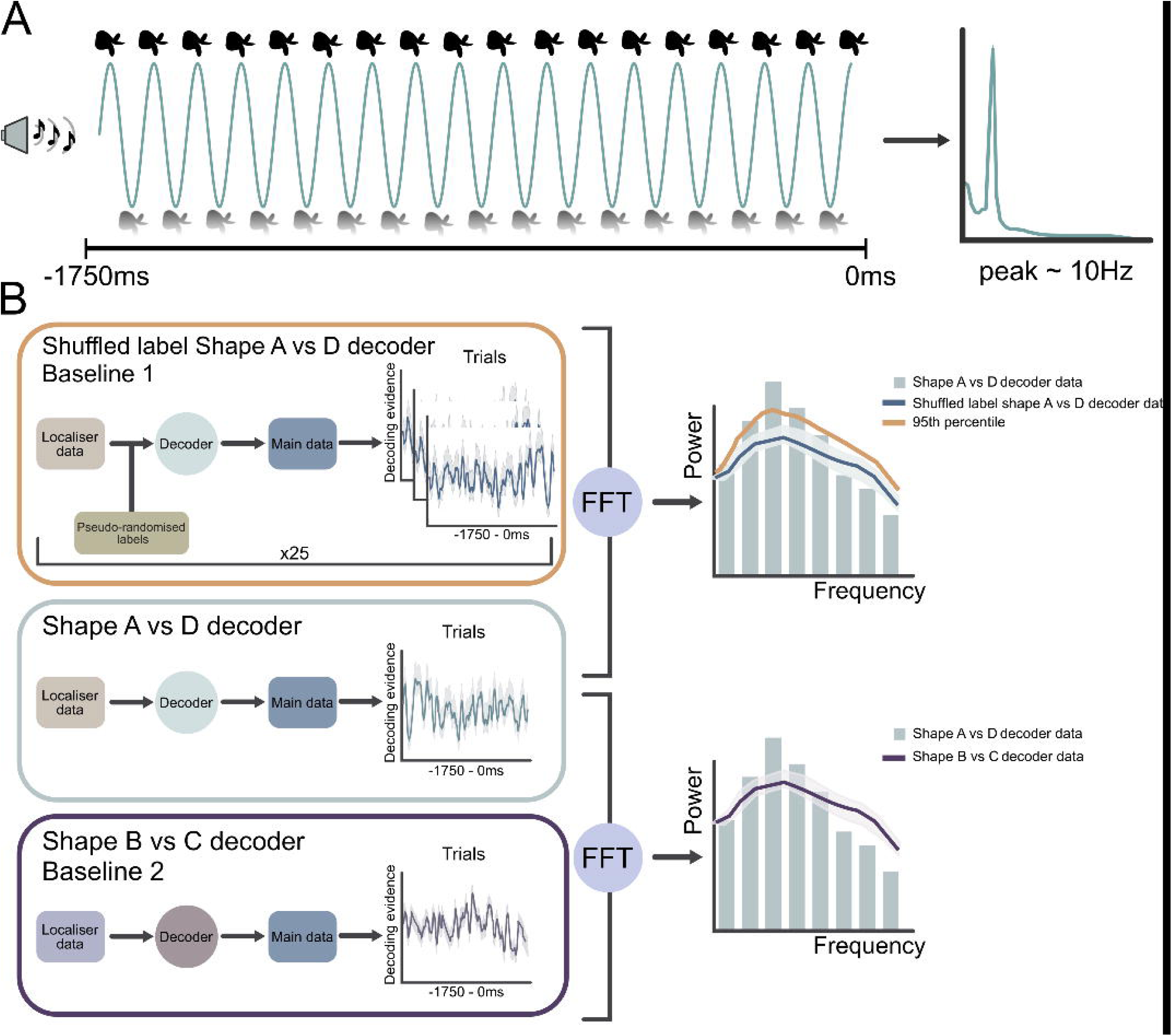
Shape prediction frequency analysis pipeline. **A:** Schematic of the hypothesis: cue-induced predictions oscillate in the alpha frequency band (∼10Hz) in the interval between predictive cue and stimulus onset (−1750 to 0ms). **B:** A decoder was trained to discriminate between shapes A and D in the localiser runs. This decoder was applied to the pre-stimulus time window in prediction runs (−1750 to 0ms). Trial-based pre-stimulus decoding time series were subjected to FFT. The resulting power spectrum was compared to the 95^th^ percentile of an empirical null distribution generated by bootstrapping decoders trained with pseudo-randomised labels (Baseline 1, top), as well as to a decoder trained on shapes only presented in the localiser (shapes B and C) (Baseline 2, bottom).

Shape decoders were trained on the localiser (70 to 200ms post-stimulus) and applied to the pre-stimulus prediction time window (−1750 to 0ms) in a time-resolved manner (sliding window of 28ms, steps of 5ms). We applied Fast-Fourier Transformation (FFT) on a single trial basis to examine the frequency attributes of the resulting pre-stimulus decoding time series (Fig. 3B). This analysis revealed that the decoded predictions oscillated at low frequencies, predominantly in the alpha frequency band (10 – 12Hz) (Fig. 4A). We identified significant power differences between the shape A vs. D decoding data and Baseline 1, specifically at 10Hz and 11Hz, exceeding the 95^th^ percentile of the empirical null distribution (both p < 0.001). It is important to note that the baseline was based on the exact same pre-stimulus data, the only difference lies in the shuffling of the shape labels for the training of the decoder. There was also a noticeable difference in the power of very low frequencies (2 – 7Hz) when comparing the shape A vs. D decoder data to Baseline 1 data. The nature of this power difference currently is not fully understood and requires further investigation. However, similar patterns have been observed in previous research^22^.

**Fig. 4:**
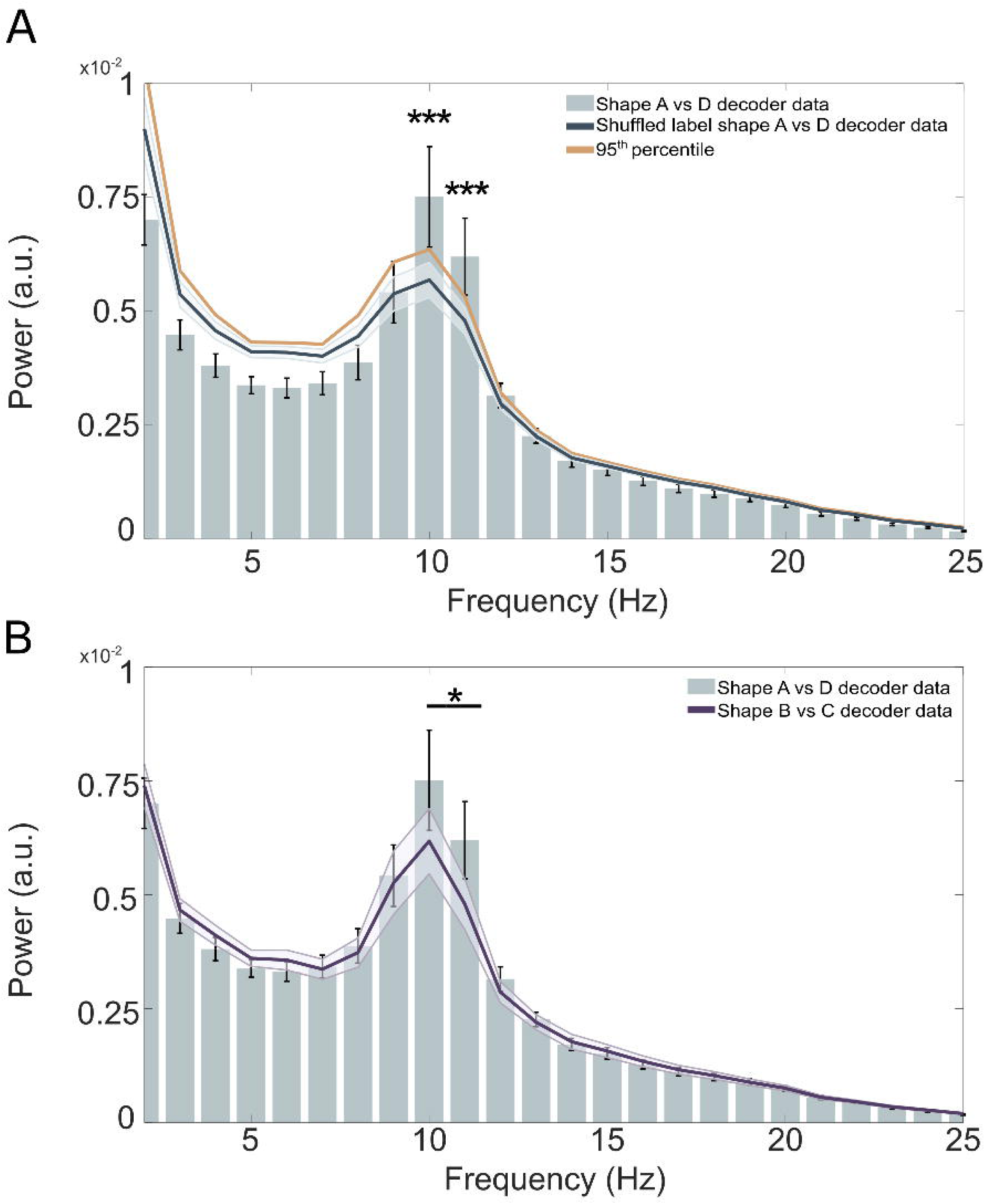
Auditory cue-induced prediction templates fluctuate at alpha frequencies. **A:** The power spectrum of pre-stimulus (−1750 to 0ms) shape decoding shows significant deviations from an empirical null distribution at 10Hz and 11Hz (***p < 0.001). The baseline power spectrum (dark blue line) was obtained by bootstrapping (n = 1000) shuffled label decoding data (n = 25 per participant). Mean and shaded regions indicate SD. Solid orange line indicates the 95^th^ percentile of the null distribution. Error bars indicate SEM. **B:** Pre-stimulus (−1750 to 0ms) MEG data shows significantly higher 10 – 11Hz power for shape A vs. D decoding than for shape B vs. C decoding (*p < 0.05). Bars indicate power of shape A vs. D decoding; dark purple line indicates power of B vs. C decoding (applied to identical pre-stimulus prediction data). Shaded regions and error bars indicate SEM.

For further validation, we also compared the shape A vs. D decoding power spectrum to the spectrum of shape B vs. C decoding (Baseline 2). Based on our initial findings, here we averaged over the 10 and 11Hz frequency bins of the two spectra. This analysis revealed significantly higher alpha power for shape A vs. D decoding than for shape B vs. C decoding in the pre-stimulus window (paired one-sided t-test, p = 0.024, t(31) = 2.067) (Fig. 4B). As before, it is important to note that the two spectra were based on the exact same pre-stimulus MEG data, the only difference lies in which shapes the decoders were trained to discriminate. If the pre-stimulus alpha fluctuations reflected more generic shape representations, this comparison would yield no significant differences. Therefore, the difference between these two decoding spectra demonstrates that these signals were highly specific to the shapes predicted by the auditory cues. It is important to note that the shape decoders were trained on signals evoked by task-irrelevant shapes during the localiser, ruling out contributions of explicit decision-making signals. These alpha power effects were also present in a control analysis designed to remove non-rhythmic signals, confirming the oscillatory nature of the decoded predictions (Fig. S2). In sum, both analyses revealed that visual predictions induced by auditory cues led to neural representations of the predicted shapes fluctuating at an alpha rhythm prior to stimulus onset.

### Predictive cues lead to improved shape discrimination accuracy

In addition to neural representations, we tested whether the predictive cues affected behavioural performance. As a reminder, participants were required to indicate whether two abstract shapes presented in succession were the same or different. It should be noted that any effects of the predictive cues on performance are not trivial, given that the shape discrimination task was orthogonal to the prediction manipulation (i.e., the cues predicted the identity of the first shape, but did not inform participants whether the two shapes would be identical or different). Still, valid predictive cues might improve performance indirectly by enhancing processing of the initial shape, facilitating discrimination of the subsequent shape^27,28^. Vice versa, invalid cues might perturb performance by impeding the processing of the initial shape. In line with this, shape discrimination accuracy was significantly influenced by whether the auditory cue correctly predicted the identity of the first shape (accuracy valid = 70% ± 1.2% and accuracy invalid = 67% ± 1.3%, mean ± SEM; t(31) = 3.215, p = 0.003; Fig. 5A). There was no difference in reaction times (valid = 614ms ± 1.3% and invalid = 615ms ± 1.3%, mean ± SEM; p = 0.626, t(31) = −0.492). Together, this suggests that valid predictions facilitated shape processing, enabling improved discrimination performance.

**Fig. 5:**
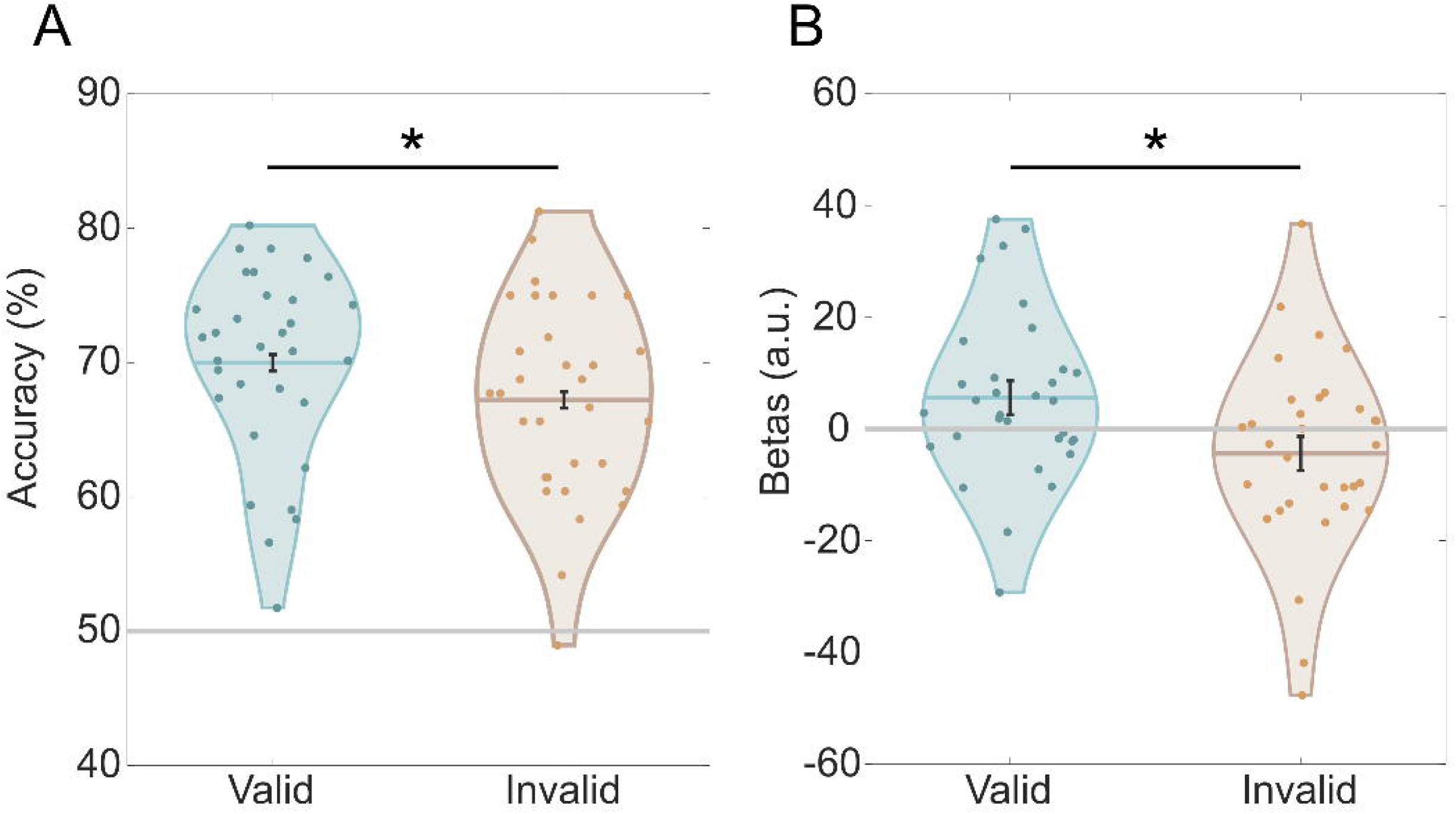
Oscillatory power of predicted shape representations modulate behavioural accuracy. **A:** Participants were able to discriminate the two presented shapes more accurately when the auditory cue validly predicted the identity of the first shape (*p < 0.05). Dots represent individual participants, error bars indicate within-participant SEM^31,32^. **B:** Output of the logistic regression (betas) between the power of pre-stimulus decoding (−500 to 0ms), averaged over 9 – 12Hz frequency bins, and discrimination performance, separately for valid and invalid prediction trials (*p < 0.05). Dots represent individual participants; error bars indicate within-participant SEM.

### Oscillatory power of predicted shape representations modulates behavioural expectation effects

If the strength of perceptual predictions indeed modulates perceptual discrimination, there should be an opposite relationship between stimulus-specific pre-stimulus oscillations and behavioural performance on valid and invalid trials. To test this hypothesis, we performed a logistic regression analysis predicting behavioural accuracy from stimulus-specific oscillatory power in the alpha frequency range (9 – 12Hz), separately on valid and invalid trials. In line with previous literature relating oscillatory power to behavioural outcome^8,29,30^, we limited the time window of interest to - 500ms to 0 pre-stimulus, since prediction signals immediately preceding stimulus onset are most likely to impact perceptual performance^28^. This analysis revealed a significant difference between valid and invalid prediction trials (paired t-test, p = 0.026, t(31) = 2.332), with a numerically positive relationship between pre-stimulus shape-specific alpha power and performance on valid trials, and a numerically negative relationship on invalid trials (Fig. 5B). Note that the individual parameter estimates for valid and invalid trials were not significantly different from zero, while the difference between the two was.

This likely reflects the fact that the individual conditions also contain non-specific trial-by-trial variance in alpha power and behavioural performance (e.g. due to fluctuations in alertness) that are subtracted out in the valid vs. invalid comparison. Importantly, the differential relationship between alpha power and behaviour dependent on prediction validity rules out any non-specific explanations of our results, and demonstrates a strong link between neural and behavioural effects of prediction. In short, pre-stimulus content-specific alpha oscillations modulated subsequent shape discrimination accuracy, such that the difference in accuracy between validly and invalidly predicted shapes was greater when pre-stimulus alpha power was higher.

### Stimulus predictions are driven by relatively late sensory representations

In an exploratory analysis, we investigated whether perceptual predictions in the alpha band reflected early or late visual representations, by dividing the training time period (70 – 200ms) into two separate windows, centring around the first (105ms) and second peak (175ms) of the localiser decoding results, respectively (Fig. 2A). These two distinct time windows also appeared to form two distinct clusters (90 – 120ms and 160 – 190ms) in the temporal generalisation matrix, with reduced cross-decoding between the two clusters suggesting qualitatively different representations (Fig. S1A). For the early training window (90 – 120ms), frequency analysis of the pre-stimulus decoding time series revealed no power differences in the alpha band (10Hz: p = 0.128; 11Hz: p = 0.062) between the shape A vs. D decoding data and an empirical null distribution (Fig. S3A). Logistic regression analyses also indicated no meaningful relationship between alpha power and behaviour on valid and invalid prediction trials (p = 0.165, t(31) = 1.423; Fig. S3B) for this training window. However, training the decoder on the later time window (160 – 190ms) revealed significantly higher pre-stimulus alpha power (10Hz: p = 0.003; 11Hz: p = 0.008) in the shape A vs. D decoding data compared to a null distribution (Fig. S3D). Logistic regression also revealed a robust difference in the relationship between pre-stimulus shape-specific alpha power and performance on valid and invalid trials (p = 0.002, t(31) = 3.359, Fig. S3E). Lastly, there was a significant difference in the average power in the 10 and 11Hz frequency bins of pre-stimulus shape A vs. D decoding between the early and late training time windows (p = 0.0103, t(31) = −2.7313). This is striking since these power spectra were calculated on the exact same MEG pre-stimulus data, the only difference was the localiser time window (90 – 120ms vs. 160 – 190ms) on which the decoder was trained. In sum, oscillating predictions seem to reflect relatively late sensory representations (160 – 190ms), rather than early feedforward-sweep-like signals.

## Discussion

The present study examined the mechanisms through which predictions exert their influence on perception. Specifically, we tested whether the content of perceptual predictions was represented in oscillations, and whether the power of this representation modulated performance on a visual discrimination task. To this end, we used multivariate decoding of MEG data to obtain the frequency spectrum of predicted shape representations. We revealed that predicted shape representations were strongest in the alpha frequency band (10 – 11Hz) (Fig. 4A-B). Furthermore, we found that this shape-specific alpha power modulated task performance, such that higher alpha power resulted in stronger expectation effects on shape discrimination (Fig. 5B). Together, these findings demonstrate that sensory templates of predicted visual stimuli are represented in the pre-stimulus alpha rhythm, which subsequently modulate performance on a perceptual discrimination task.

Previous studies have hypothesised that oscillations play a critical role in conveying perceptual predictions^7,8,12,19,30^. This is largely based on indirect evidence, consisting of a range of studies finding that pre-stimulus alpha oscillations modulate performance on perceptual discrimination tasks^12,15,16,29^. Further, there is a second body of evidence that links experimental manipulations regarding stimulus predictability to the power of low frequency oscillatory activities^8,20,21^. Finally, a recent study has demonstrated a link between pre-stimulus high alpha/low beta power and the occurrence of high confidence false percepts^33^. However, the key hypothesis that neural oscillations actually convey the contents of perceptual predictions has remained largely untested. In the current study, we present evidence that the content of predicted shapes is represented in pre-stimulus alpha oscillations, providing direct support that perceptual predictions are indeed conveyed through neural oscillations.

While the role of pre-stimulus alpha oscillations has been extensively studied, it remains controversial. Previous studies have reported alpha oscillations typically being stronger when visual stimuli are not present, or actively not-attended^10,34,35^. This has led to the hypothesis that alpha is predominantly an inhibitory rhythm. However, our results demonstrate that alpha oscillations are not solely inhibitory, but play an active role in conveying prior knowledge. The link between pre-stimulus alpha power and expectation effects on perception revealed in the current study suggests that whether alpha facilitates or inhibits sensory processing depends on whether inputs match or mismatch current predictions.

Predictive processing theories of perception highlight a key role for prior predictions in guiding inference in the brain^4,36^. While there is convergent evidence that the brain contains predictive signals^8,27,37,38^, the mechanisms through which the brain deploys these predictions remains largely unclear. Predictive coding has been suggested to involve rhythmic interactions between different frequency band activities^1,5^, where high frequency gamma is responsible for feedforward signalling (originating predominantly from superficial layers) and alpha/beta oscillations exert top-down control (feedback predictions), emerging from deep cortical layers. Indeed, animal work investigating the frequency characteristics and cortical layer specificity of predictable information processing^6,39^ revealed that pre-stimulus alpha power is an indicator of stimulus predictability, originating from cortical layers involved in feedback signalling^6^. Our results extend these intracranial electrophysiological observations by relating pre-stimulus alpha oscillations to the contents of feedback signalling.

Exploratory analyses revealed that the oscillating prediction signals reflected relatively late sensory representations (160 – 190ms localiser training window, Fig. S3D). We speculate that during this time period, the sensory representations captured by the decoder reflected an integration of bottom-up inputs and top-down recurrence, rather than solely the first feedforward sweep. Like the current study, previous studies have also revealed top-down modulations that reflected relatively late post-stimulus representations (i.e., 120 – 200ms)^28,40^. This may explain why predictions have been shown to modulate later sensory processing, while leaving the early feedforward sweep (< 80ms post-stimulus) mostly untouched^37,41^.

Rather than predictions being actively conveyed in an alpha rhythm, an alternative explanation of our results may be that prediction signals passively ride on ongoing alpha oscillations. Alpha oscillations are the most prominent frequency band in the awake human brain, especially in the visual cortex, and even a non-oscillatory top-down signal arriving in visual cortex may inherit these alpha rhythms. Given our finding that shape-specific alpha power has opposite effects on behaviour dependent on the validity of the predictions, such a more passive explanation seems less likely. However, future research is indeed needed to properly distinguish between these hypotheses.

In addition to alpha power, alpha phase has also commonly been reported as influencing perception^14,17–19^. Specifically, the phase of ongoing alpha oscillations has been suggested to modulate perception by creating optimal and suboptimal periods visual processing through top-down control^42^. Combining this with the role of predictions in perception, one might hypothesise that the brain switches between sensing (bottom-up) and predicting (top-down) at opposite phases of alpha oscillations. Conceptually in line with this idea, Weilnhammer et al. demonstrated that the brain indeed switches between externally and internally biased perceptual modes^43^, albeit at a slower timescale. By analysing perceptual decision-making in humans and mice, this study revealed fluctuations of enhanced and reduced sensitivity to external stimuli. When sensitivity was low, the brain tended to depend more on perceptual history of the learnt sequences of stimulus presentation (i.e., prior knowledge). Our results showing predicted stimulus content fluctuating in the alpha rhythm are potentially in line with this proposal. Specifically, representations of predicted shapes would be hypothesised to dominate at the alpha phase optimised for predicting, and be absent at the phase optimised for bottom-up processing (Fig. 3A). Based on this, the brain may use alpha oscillations as neural mechanism to balance perceiving and predicting, where not only power but also phase has a crucial role to play. The current study was not optimised to test this hypothesis, since the shape discrimination task was orthogonal to the prediction cues. Future work directly probing the effect of predictions on subjective perception^37,44,45^ should test this hypothesis by relating alpha phase to expectation effects on perception.

Many prominent and influential theoretical frameworks have long speculated on the role of neural oscillations in perception^10,11,19^. Here we shed light on this by showing that alpha oscillations convey perceptual predictions, and modulate subsequent perceptual performance. These findings enrich current models of perceptual inference in the human brain by revealing the neural mechanisms through which predictions are kept online in order to guide perception.

## Methods

### Participants

Sixty-two healthy right-handed participants (43 female) with normal or corrected-to-normal vision and no history of neurological disorders took part in the behavioural experiment. This experiment served as a pre-assessment process to familiarise the participants with the task and select only those whose average performance accuracy on the challenging shape discrimination task was above 70% across the four runs. Thirty-nine participants (28 female) met the performance inclusion criteria and participated in the MEG experiment. Seven participants were excluded from subsequent analyses due to excessive head movement (N=5) or not completing the full experiment (N=2), leaving thirty-two participants (23 female, age 26 ± 5 years, mean ± SD) for the MEG analysis.

### Stimuli

The experiment employed the same design as Kok & Turk-Browne^46^, wherein participants discriminated between two consecutively presented shapes which were preceded by a predictive auditory cue. Each predictive cue was composed of three pure tones (440, 554, and 659Hz; 80ms per tones; 5ms intervals), played with rising or falling pitch, with a total duration of 250ms. Visual stimuli were generated using MATLAB (The MathWorks Inc., version 2021b) and Psychophysics Toolbox^47^. The visual stimuli consisted of complex abstract shapes defined by radial frequency components (RFCs)^48^. To define the contours of the stimuli, seven RFCs (0.55Hz, 1.11Hz, 4.94Hz, 3.39Hz, 1.54Hz, 3.18Hz, 0.57Hz) were used which were based on a subset of stimuli from Op de Beeck et al.’s work^49^; see their Fig. 1A). The amplitudes of two RFCs (1.11Hz, and 1.54Hz components) were varied to create a two-dimensional shape space. Specifically, four shapes were created such that discrimination between shapes A (amplitude of 1.11Hz = 8; 1.54Hz = 8) and D (amplitude of 1.11Hz = 26; 1.54Hz = 26) was orthogonal to discrimination between shapes B (amplitude of 1.11Hz = 8; 1.54Hz = 26) and C (amplitude of 1.11Hz = 26; 1.54Hz = 8) (Fig. 1D). Additionally, RFC-based warping was used to generate moderately distorted versions of the two main experiment shapes (shape A and D, Fig. 1D) for the benefit of the shape discrimination task. This warp to define the shape was achieved by modulating a different RFC’s amplitude (3.18Hz) than the two used (1.11Hz and 1.54Hz) to define the shape space. This modulation could be either positive or negative (counterbalanced over conditions) and was orthogonal to the shape space used for the two main experiment shapes, and therefore to the cue predictions as well. The visual stimuli were displayed on a rear-projection screen using a projector (1024 x 768 resolution, 60 Hz refresh rate) against a uniform grey background.

### Behavioural experiment

The study had two parts, a behavioural training and screening experiment, and an MEG experiment for those who passed the behavioural screening. In both parts, participants were engaged in a shape discrimination task. Each trial started with a fixation bullseye (diameter, 0.7°) for 100ms, followed by the presentation of two consecutive shape stimuli each for 250ms, and separated by a 500ms blank screen containing only a fixation bullseye (Fig. 1A). On each trial, the second shape was the same as the first or slightly warped. The modulation was either positive or negative, and the size of the modulation was determined by an adaptive staircasing procedure^50^, updated after each trial, in order to make the task challenging. Participants were instructed to report whether the two presented shapes were identical or different. After the response interval ended (750ms after disappearance of the second shape), the fixation bullseye was replaced by a single dot, signalling the end of the trial while still prompting participants to fixate. On each trial, one of the four shapes (A, B, C or D; Fig. 1D) was presented, in a counterbalanced (i.e., non-predictable) manner. Participants performed four runs (360 trials in total) of the shape discrimination task, maximum one week prior to the MEG session.

### MEG experiment

The MEG experiment started with two localiser runs, containing the same four abstract shapes as in the behavioural task. To ensure participants were engaged, they performed a fixation dimming task (10% of total trials, ∼24 of 248 trials per run). Each trial began with a fixation bullseye (visual angle: 0.7°) displayed for 100ms, followed by one of the four shapes presentation for 250ms. Following the stimulus presentation, the fixation bullseye reappeared and remained on the screen for a period between 1000 and 1200ms. In 10% of the trials, fixation bullseye dimmed for 150ms and participants had been instructed to press a button when this occurred. By using identical stimulus durations, these runs were designed to be as similar as possible in terms of stimulus presentation to the main experiment. During the localisers, participants correctly detected 95.3 ± 0.7% (mean ± SEM) of fixation dimming events and incorrectly pressed the button on 4.9 ± 2.2% of trials, suggesting that participants were successfully engaged by the fixation task.

Following the localiser runs, participants performed 8 main task runs (2x training runs, 6x prediction runs), 64 trials per run, in total 512 trials. During the prediction runs, an auditory cue (falling vs. rising tones, 250ms) was presented 100ms after trial onset. Following a 1500ms interval, two consecutive shape stimuli were displayed (each for 250ms) and, separated by a 500ms blank screen (Fig. 1A). The auditory cue predicted whether the first shape presented on that trial would be shape A or D. The cue was valid on 75% of trials, whereas on the other 25% of trials the unpredicted shape would be presented (Fig. 1B). For instance, if the cue was a falling auditory tone, it might lead to shape A in 75% of cases and shape D in the other 25% of cases. Note that shapes B and C were never presented in the prediction runs. The contingencies between cues and shapes were flipped halfway through the experiment, and the order was counterbalanced over subjects. Prior to the first prediction run, and after the cue reversal halfway through, participants were trained on the cue–shape associations during training runs in the MEG and explicitly informed about the cue contingencies. In the training runs, the auditory cue was 100% predictive of the identity of the first shape.

### Pre-processing

Whole-head neural recordings were obtained using a 273-channel MEG system with axial gradiometers (CTF Systems) at a rate of 600Hz located in a magnetically shielded room. Throughout the experiment, head position was monitored online and corrected if necessary using three fiducial coils that were placed on the nasion, right and left preauricular. If participants moved their head more than 5mm from the starting position, they were repositioned after each run. Eye movements were recorded using an EyeLink 1000 infrared tracker (SR Research Ltd.). The recorded eye-tracker data were used to identify eye-blink related artefacts in the MEG signal. Auditory tones were delivered using earplugs (Etymotic Research Inc.). A photodiode was placed at the bottom left corner of the screen to measure visual stimulus presentation latencies. The photodiode signal was used to realign the MEG signal with stimulus onset.

The data were pre-processed offline using FieldTrip ^51^. The variance (collapsed over channels and time) was calculated for each trial in order to identify artefacts. Trials with large variances were subsequently selected for manual inspection and removed if they contained excessive and irregular artefacts. Next, independent component analysis was used to further remove cardiac and eye movements related artefacts. The independent components were correlated to the eye tracking signal to identify potentially contaminating components for each participant, and inspected manually before removal. For the main analyses, data were high-pass filtered using a two-pass Butterworth filter with a filter order of five and a frequency cut-off of 0.1Hz. Notch filters were also applied at 50, 100, and 150Hz to remove line noise and its harmonics. No detrending was applied for any analysis. Finally, main task data were baseline corrected on the interval of −200 to 0ms relative to auditory cue onset, and localiser data were baseline corrected on the interval of −200 to 0ms relative to shape onset.

### Decoding analysis

To reveal the representational content of neural activity, a decoding analysis was applied. We used an LDA decoder^24^, which described how activity at the sensor-level varied as a function of a discriminability index. Unlike conventional LDA which separates data into discrete categories, our customised decoder calculated the distances of each test sample to the hyperplane, treating these distances as discriminant evidences. Thereby, we obtained a continuous measure of which shape was encoded in the neural signals, providing finer resolution in analysing the neural representations. The decoding analysis was performed in a time-resolved manner by applying it sequentially at each time point, in steps of 5ms and averaging over a 28ms time window centred at that specific time point. Thereby, the decoder effectively down-sampled the data (from 600Hz original sampling rate) to 200Hz.

To test how effective the decoder was at revealing neural patterns, it was first trained and tested on shape A and D trials (between −100 and 600ms relative to stimulus onset) from the localiser runs, using a leave-one-block-out approach. Analogously, a shape B vs. C decoder was tested on shape B and C trials. To further validate the analysis, we tested the shape A vs. D decoder on the shape B and C trials, and the shape B vs. C decoder on shape A and D trials (Fig. S1). We expected significant decoding within shape categories (e.g. training and testing on shape A vs. D), but not across shape categories (i.e., training on shape A and D and testing on shape B and C, and vice versa).

Localiser decoding results were analysed using non-parametric cluster-based permutation tests. The data were represented as 2D matrices of decoding performance, with training time on one axis and testing time on the other. The statistical analysis focused on identifying significant 2D clusters in these matrices. To do so, univariate t-statistics were calculated for the entire matrix. Elements that were considered neighbours, i.e., directly adjacent in cardinal or diagonal directions, were collected into separate positive and negative clusters if they passed a threshold corresponding to a p-value of 0.001 (two-tailed). The significance of the clusters was assessed by summing the t-values within each cluster to obtain cluster-level test statistics. These test statistics were then compared to a null distribution, which was created by randomly shuffling the observed data 10,000 times. A cluster was considered significant if its resulting p-value was less than 0.05 (two-tailed).

In order to reveal predicted shape representations, the decoder was trained on shape A vs. D localiser trials (70 – 200ms), and subsequently tested on the pre-stimulus window (−1750 to 0ms relative to shape onset) during the prediction runs. To address label imbalances resulting from trial rejections during pre-processing, random resampling was applied to the training sets, ensuring an equal number of each decoded classes (shapes) for every participant. Furthermore, we repeated the same procedure for each participant using a control decoder trained on shapes B vs. C localiser trials, i.e. shapes which were not presented during the prediction runs. This results of applying this control decoder to the pre-stimulus prediction window served as a baseline (Baseline 2, Fig. 3B - bottom) in further analyses. It is important to highlight that the shape B vs. C discrimination was orthogonal to shape A vs. D discrimination.

### Frequency analysis of pre-stimulus decoding time series

Our primary aim was to test whether the decoded neural representations of predictions had oscillatory dynamics. Therefore, we adapted the analysis approach of Kerrén et al.^22^, investigating the frequency characteristics of decoder time series using FFT. This analysis was applied to pre-stimulus decoding time courses (−1750ms until 0ms relative to stimulus onset), based on the averaged decoder training time window of 70 – 200ms. We chose this training time window based on the results of localiser decoding. In an exploratory analysis, we repeated the analysis for two shorter training time windows (90 – 120ms and 160 – 190ms), centred around the first (105ms) and second peak (175ms) of localiser shape decoding (Fig. 2A-B) to distinguish effects of earlier and later representations. These time windows were chosen since they appeared to form distinct clusters in the localiser decoding temporal generalisation matrix, with reduced cross-decoding between the two clusters suggesting qualitatively different representations (Fig. S1A). For each participant, each trial of the pre-stimulus decoded time series was tapered with a Hann window covering the whole time period (−1750 to 0ms), and then subjected to the FFT. In a control analysis, we used Fitting Oscillations and One-Over-F (FOOOF, as implemented in the Fieldtrip toolbox^52^), which separates rhythmic activity from concurrent power-spectral 1/f modulations in electrophysiological data, to validate the oscillatory nature of the predictive representations.

To assess the reliability of our results, we created an empirical baseline using decoders with randomly shuffled shape labels (Baseline 1, Fig. 3B - top). The labels of the two shapes (shape A and D) were shuffled pseudo-randomly before training the decoder, 25 times per participant. Therefore, each participant yielded 25 permuted datasets. The analysis parameters for the baseline decoding were identical to the non-shuffled decoder, i.e. identical spectral analysis was performed for each of the 25 datasets per participant. We generated an empirical null distribution using bootstrapping of the permuted datasets (n = 1000)^53^, and compared this to the frequency analysis results of the non-shuffled shape A vs. D decoder data^22^. Frequency bins with higher power than the empirical null distribution (exceeding the 95^th^ percentile) were considered significant. To further validate the findings, we also conducted the identical frequency analysis (same analytical parameters) using shape B vs. C decoding time series as an additional baseline (Baseline 2, Fig. 3B - bottom).

### Relating behavioural and neural effects

To investigate whether there was a relationship between stimulus-specific pre-stimulus alpha power and shape discrimination performance, we performed a logistic regression analysis separately for valid and invalid prediction trials. Based on the existing literature relating pre-stimulus oscillatory power and phase to behavioural performance^8,29,30^, we limited the pre-stimulus decoding time series to - 500ms to 0ms relative to stimulus onset. To be able to accurately estimate pre-stimulus alpha power, yet be as close as possible to stimulus onset, we used a 500ms Hann window over the −500ms to 0ms time window, resulting in ∼2Hz frequency resolution (alpha frequency bins: 9.375Hz, 10.937Hz, 12.500Hz). Separately for valid and invalid prediction trials, trial-based power estimates of the pre-stimulus (−500ms to 0ms) alpha activity were averaged over for the three alpha frequency bins. We balanced the trial numbers by randomly choosing a subset of trials from the conditions with higher trial counts (i.e., valid). The dependent variable of the model was the behavioural outcome (correct or incorrect response), sorted separately again for valid and invalid predictions. The model parameter estimates (i.e., beta values) served as an indication of an underlying link between stimulus-specific alpha power and behavioural performance. The valid and invalid condition beta values were statistically compared using a paired t-test.

### Source localisation of shape decoding

To visualise the underlying neural sources during decoding, we applied source localisation analyses using an LCMV beamformer^25^. The spatial distribution of the underlying signal during classification in LDA is primarily influenced by the magnetic field difference between the two experimental conditions. Therefore, one can visualise the source of a decoder by estimating the sources of the two different conditions, and compute the difference^26^. Based on previous studies demonstrating that the anatomical specificity gain of using subject-specific anatomical images is negligible^54^, we did not collect individual anatomical MRI scans for our subjects. We followed a group-based template approach using a template MRI (in MNI space) in combination with a single shell head model and a standard volumetric grid (8mm resolution), as present in the Fieldtrip toolbox. Participants’ individual fiducials were used to generate a participant-specific forward model in MNI space. The spatial filter was computed for the time window of interest (70 – 200ms, decoder training window) in the averaged data, which was subsequently applied separately to the two conditions of interest (valid and invalid prediction trials). For shape A vs. D decoding a percentage absolute signal change was computed in source space, to determine which source signals were involved in discriminating between shape A and D without making assumptions about the sign of the dipole.

## Acknowledgements

This work was supported by a Wellcome/Royal Society Sir Henry Dale Fellowship [218535/Z/19/Z] and a European Research Council (ERC) Starting Grant [948548] to P.K. The Wellcome Centre for Human Neuroimaging is supported by core funding from the Wellcome Trust [203147/Z/16/Z].

## Conflict of interests

The authors declare no conflicts of interests.

---

**Fig. S1:**
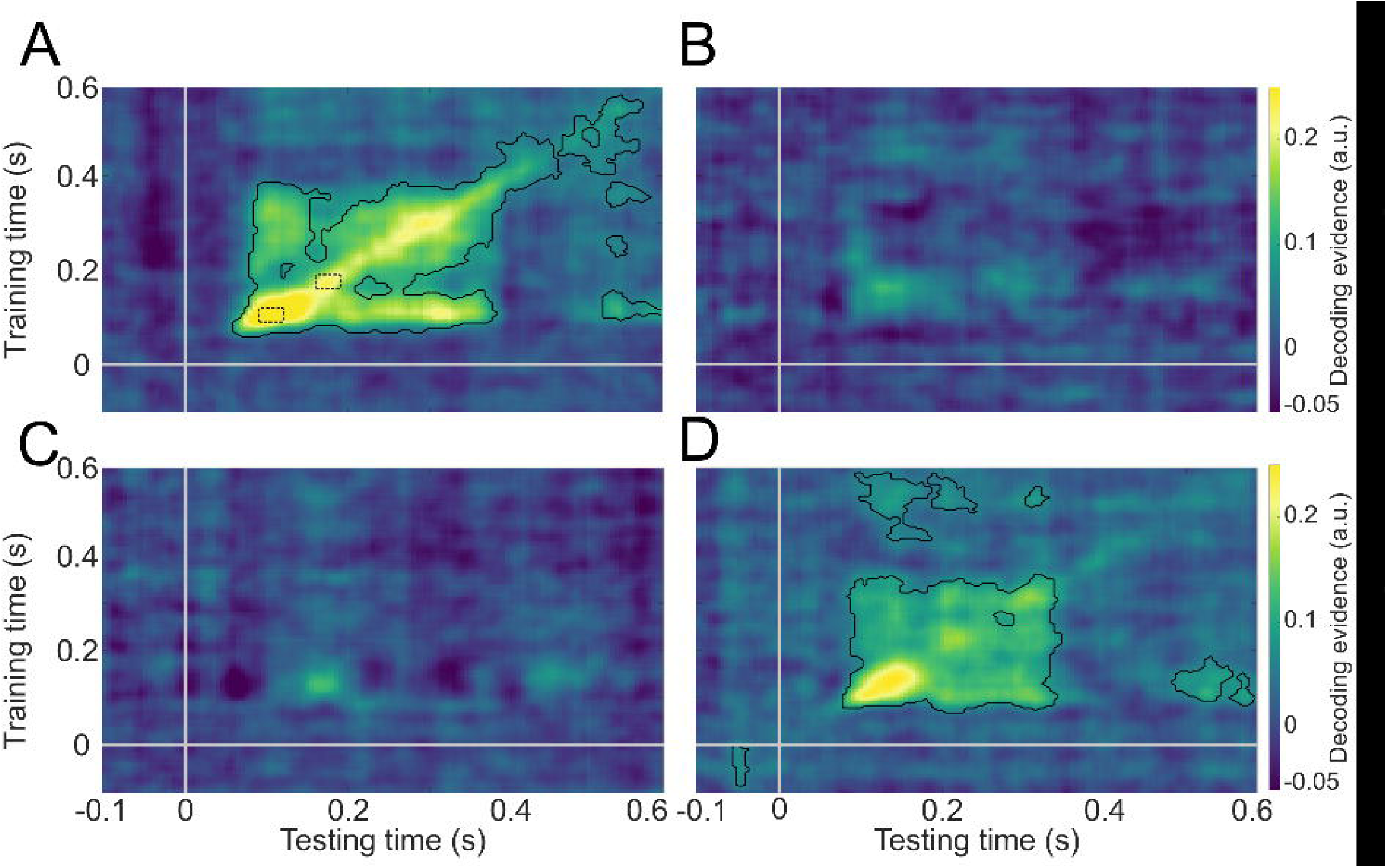
Localiser temporal generalisation of shape A vs. D and shape B vs. C decoders. **A:** Shape A vs. D decoder, trained and tested (−100 to 600ms) on the two target shapes (Shape A and D) which appeared both in the localiser and the prediction runs. Dashed rectangles indicate clusters used in exploratory early vs. late analysis (90 – 120ms and 160 – 190ms, respectively). **B:** Shape B vs. C decoder tested on Shape A and D trials. No significant clusters were identified. **C:** Shape A vs. D decoder tested on Shape B and C trials. No significant clusters were identified. **D:** Shape B vs. C decoder, trained and tested on Shapes B and C. Solid black lines indicate significant clusters (p < 0.05). Solid grey lines at 0ms indicate stimulus onset.

**Fig. S2:**
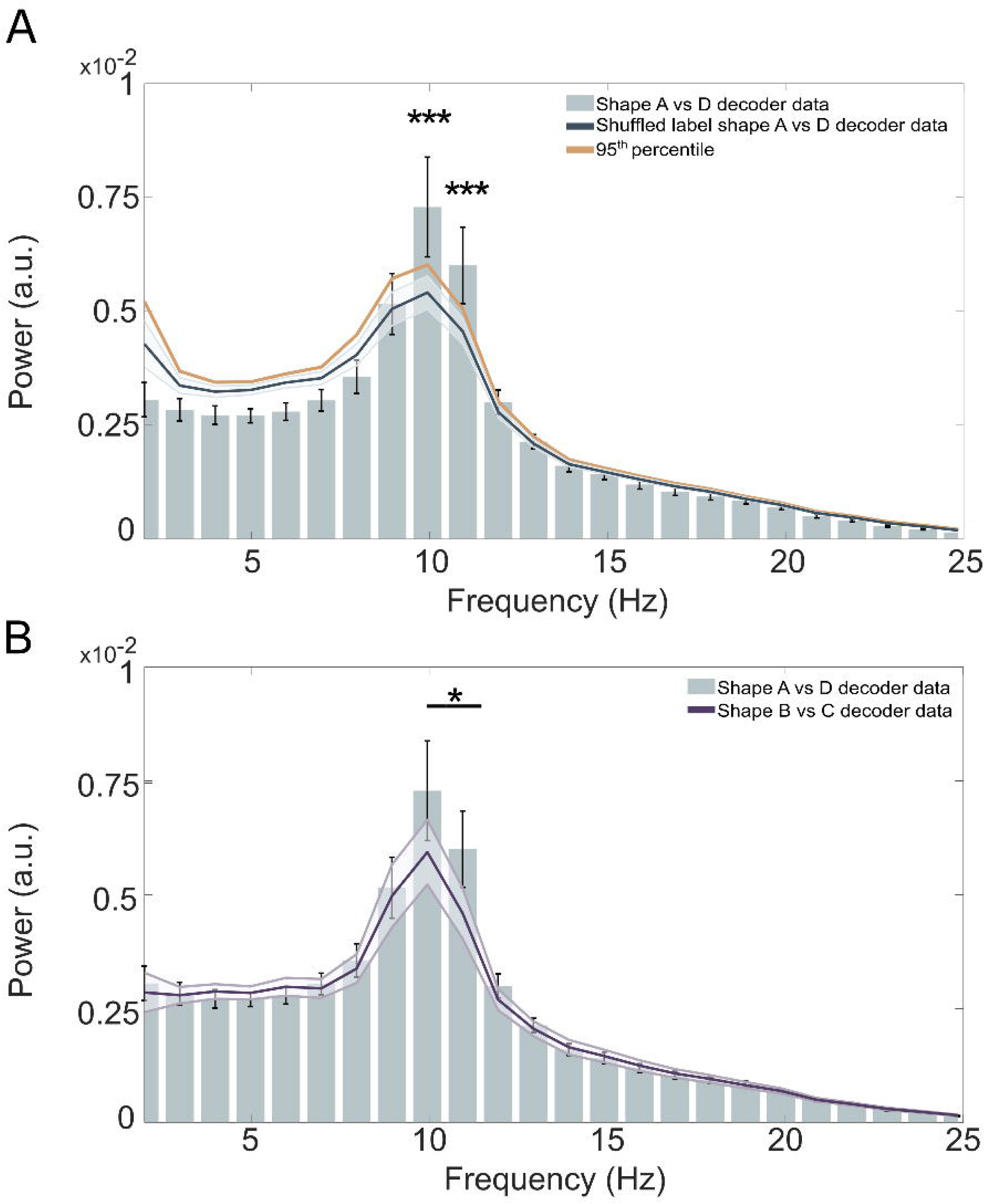
Prediction templates fluctuate at alpha frequencies – 1/f noise removed. **A:** Significant power differences at 10Hz and 11Hz between Shape A vs. D decoding data and Baseline 1 after the subtraction of 1/f noise (***p < 0.001). This further demonstrates that the stimulus-specific alpha signals likely reflect oscillations. The baseline power spectrum (dark blue line) was obtained by bootstrapping (n = 1000) the randomly shuffled label decoding data (n = 25 per participant). Mean and shaded regions indicate SD. Solid orange line indicates the 95^th^ percentile of the generated baseline distribution. Error bars indicate SEM. **B:** Pre-stimulus (−1750 to 0ms) MEG data after 1/f removal shows significantly higher 10 – 11Hz power for shape A vs. D decoding than for shape B vs. C decoding (*p = 0.022, t(31) = 2.107). Dark purple line indicates the power spectrum of B vs. C decoder (tested on the identical pre-stimulus main experiment prediction data (as shape A and D). Mean and shaded regions indicate SEM. Error bars indicate SEM.

**Fig. S3:**
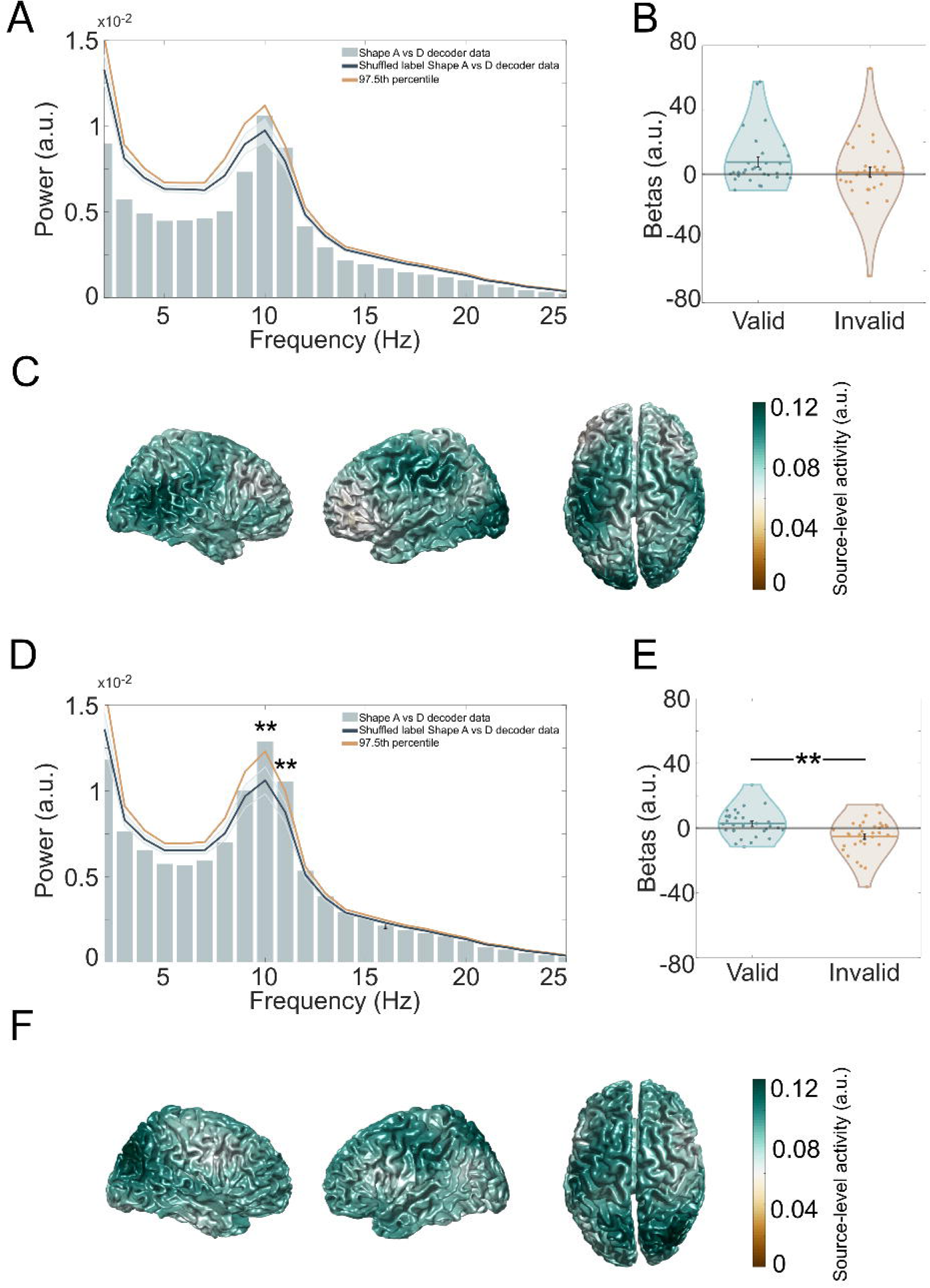
Late sensory representations drive stimulus predictions. **A:** Power spectrum of the −1750 to 0ms prediction time window shape A vs. D decoding, trained on the 90 to 120ms post-stimulus localiser window. No significant distinctions between the shape A vs. D decoding data and an empirical null distribution at 10Hz and 11Hz. Mean and shaded regions indicate SD. Dark solid orange line indicates the 97.5^th^ percentile of the null distribution, implementing a one-sided test at p < 0.05 while correcting for the two time windows tested here. **B:** Output of the logistic regression (betas) between the power of pre-stimulus decoding time window of −500 to 0ms, averaged over 9 – 12Hz frequency bins, and discrimination performance, separately for valid and invalid prediction trials. No significant difference between valid and invalid prediction betas. Dots represent individual participant; error bars reflect within-participant SEM. **C:** The difference in source localisation for shape A and D during the localiser training time window of 90 – 120ms post-stimulus. **D:** Power spectrum of the −1750 to 0ms prediction time window shape A vs. D decoding, trained on the 160 to 190ms post-stimulus localiser window. Statistically significant difference from an empirical null distribution at 10Hz and 11 Hz (**p< 0.01). The baseline power spectrum (dark blue line) was calculated as before. Mean and shaded regions indicate SD. Solid orange line indicates the 97.5^th^ percentile of the baseline distribution. **E:** Logistic regression between pre-stimulus decoding power (−500 to 0ms, averaged over 9 – 12Hz frequency bins) and discrimination performance between valid and invalid prediction trials (**p < 0.01). Dots represents individual participants; error bars were calculated as within-participant SEM. **F:** The difference in source localisation for shape A and D during the localiser training time window of 160 – 190ms post-stimulus.

## Notes

### Competing Interest Statement

The authors have declared no competing interest.

